# Investigation of TRMT61B methyltransferase activity on mRNA and its effects on translation

**DOI:** 10.64898/2025.12.08.692954

**Authors:** Dorthy Fang, John M. Babich, Ryan Stanton, Isaac W. Vock, Kyrillos Abdallah, Mingyi Zhu, Richard Li, Matthew D. Simon, Wendy V. Gilbert, Sigrid Nachtergaele

**Affiliations:** Department of Molecular, Cellular, and Developmental Biology, Yale University, New Haven, CT, USA; Department of Molecular Biophysics & Biochemistry, Yale University, New Haven, CT, USA; Department of Ecology and Evolutionary Biology, Yale University, New Haven, CT, USA

## Abstract

Despite recent advances in technology to map RNA chemical modifications transcriptome-wide, the distribution of N¹-methyladenosine (m¹A) in mRNA remains contested, hindering a clear understanding of its function. Additionally, the enzyme(s) that installs the majority of reported mRNA m^1^A sites has yet to be identified. In this study, we characterized TRMT61B, an m^1^A methyltransferase known to methylate mitochondrial RNAs, but whose sequence preferences have been underexplored. By integrating cellular overexpression of TRMT61B and *in vitro* methylation of a synthetic pool of diverse human RNA sequences, we identified a preference for a YMR<u>A</u> consensus motif in single stranded RNA regions. In these experiments, TRMT61B methylated thousands of novel human mRNA sites, revealing activity on cytosolic mRNAs. We used these novel m^1^A-modifiable sequences to test the effects of m^1^A on translation of luciferase reporters and on ribosome recruitment to modified transcripts in the pool. We found that m^1^A addition can significantly affect translation and ribosome recruitment, but that these effects are vary by transcript. Taken together, our results can inform future studies of TRMT61B and mRNA, and emphasize that studies of m^1^A regulation of mRNA must be carried out and interpreted in a highly context-aware manner.

## INTRODUCTION

RNA chemical modifications can profoundly affect RNA structure, metabolism, and function, and recent studies have underscored their impacts on gene expression [1]. Over 170 RNA modifications have been reported to date [2], and we are only beginning to understand their functions and mechanisms of action. *N*^1^-methyladenosine (m^1^A) is a modification that has been found in a wide range of RNA species, including tRNA, rRNA, and more recently, mRNA and long noncoding RNA (lncRNA) [3–9]. Several studies of m^1^A in mRNA suggest that it is stress-responsive and may affect mRNA stability and translation [3–6,10,11]. However, discrepancies between studies as to the number and location of mRNA m^1^A modification sites has led to significant challenges in uncovering its regulation and function in mRNA [12,13]. Across published datasets, estimates for the number of m^1^A sites have ranged from as low as fifteen to as high as several thousand, with less than ten consensus sites across these studies [3–6,14–16]. In part, this may be due to the generally lower stoichiometry (percent of copies of a transcript carrying a modification) of mRNA modification sites as compared to ncRNAs such as tRNA and rRNA. Combined with technical limitations in detection methods, variations in analytical pipelines and thresholds, and biological heterogeneity, this has led to the wide discrepancies reported.

A key step towards understanding the functions of many RNA modifications is to identify the enzymes that regulate their activity. In eukaryotes, multiple enzymes have been reported to possess m^1^A methyltransferase activity. For instance, the TRMT6/TRMT61A complex and TRMT10B are responsible for cytosolic tRNA methylation [17,18], while TRMT61B has been shown to methylate mitochondrial tRNA and rRNA (mt-tRNA and mt-rRNA) [8,9], and TRMT10C has reported activity on mt-tRNA [19]. In addition, mitochondrial mRNAs are m^1^A-methylated by TRMT10C and TRMT61B [3,20]. In nuclear-encoded mRNA, a small subset of sites have been attributed to TRMT6/TRMT61A activity on a GUUCRA (R is a purine) consensus motif in a tRNA T-loop-like structure [3,20]. However, many reported m^1^A modification sites in nuclear-encoded mRNAs do not contain this motif, suggesting that other m^1^A methyltransferases might be responsible for the modification on these mRNA transcripts.

We became particularly interested in TRMT61B, the only m^1^A methyltransferase that has reported activity on three major RNA classes: rRNA, tRNA, and mRNA. Known TRMT61B targets include several mt-tRNAs, a highly conserved site in 16*S* rRNA, and a handful of mt-mRNAs (*e.g. COX1*, *COX3*) [8,9,20]. Its diverse RNA substrates suggest more flexibility than TRMT6/TRMT61A, but its sequence and structure requirements for methyltransferase activity have not been characterized in detail [21]. This broad range of possible TRMT61B targets is particularly intriguing in light of published work that TRMT61B expression at the RNA and protein level is elevated in a wide range of cancer types, and that its expression level is positively correlated in several cancers with relevant clinical factors such as aneuploidy and increased tumor grade [22]. As has been observed for many other proteins, including RNA binding proteins, elevated expression levels and other disease-induced cellular changes could alter the subcellular distribution of TRMT61B, and thus increase the probability that it could methylate additional targets beyond the reported mitochondrial RNAs [23,24].

Motivated by the possibility that elevated TRMT61B could promote tumor growth and by the multiple classes of RNA methylated by this enzyme, we sought to further characterize potential TRMT61B targets. To achieve this, we performed both overexpression experiments with TRMT61B in cells and *in vitro* biochemical approaches to identify novel putative TRMT61B targets and to better define the sequence and structural preferences of this enzyme. We identified a YMRA (Y: cytidine or uridine, M: adenosine or cytidine, R: adenosine or guanosine, and A: the m^1^A site) consensus motif that was present in the majority of sites from both cellular and *in vitro* experiments and used individual oligonucleotides to confirm this sequence preference. Structural predictions revealed that the m^1^A methylated nucleotide was consistently predicted to sit in a single stranded RNA loop. We then tested our newly discovered TRMT61B substrates in translation and ribosome recruitment assays, and found that m^1^A has context-specific effects on these processes. Throughout this work, we used mass spectrometry and sequencing-based validation at every step to ensure that we were measuring the effects of m^1^A methylation at specific sites. Collectively, our work identifies preferred sequence and structural constraints for TRMT61B-mediated m^1^A installation on mRNA and measures the downstream effects of m^1^A methylation on translation and ribosome recruitment. These findings have implications for understanding the consequences of altered TRMT61B expression in disease, and more broadly reveal how m^1^A could influence gene expression regulation in a context-specific manner.

## RESULTS

### TRMT61B overexpression increases m^1^A modification of mRNA in cells

To explore the possibility that TRMT61B overexpression may induce m^1^A methylation of additional mRNA targets, we overexpressed TRMT61B-FLAG or a control vector in U-2 OS cells. We then incubated unfragmented, polyA-selected RNA from these cells with an anti-m^1^A antibody, and sequenced m^1^A antibody-bound RNA (m^1^A-IP) and paired input samples (Fig. 1A, Supp. Fig. 1A,B). Though concerns have been raised about the specificity of antibodies for RNA modifications, we used an anti-m^1^A antibody that we have previously characterized in detail by analyzing antibody-enriched fragmented RNA by liquid chromatography coupled to tandem mass spectrometry (LC-MS/MS) [25].

**Figure 1.**
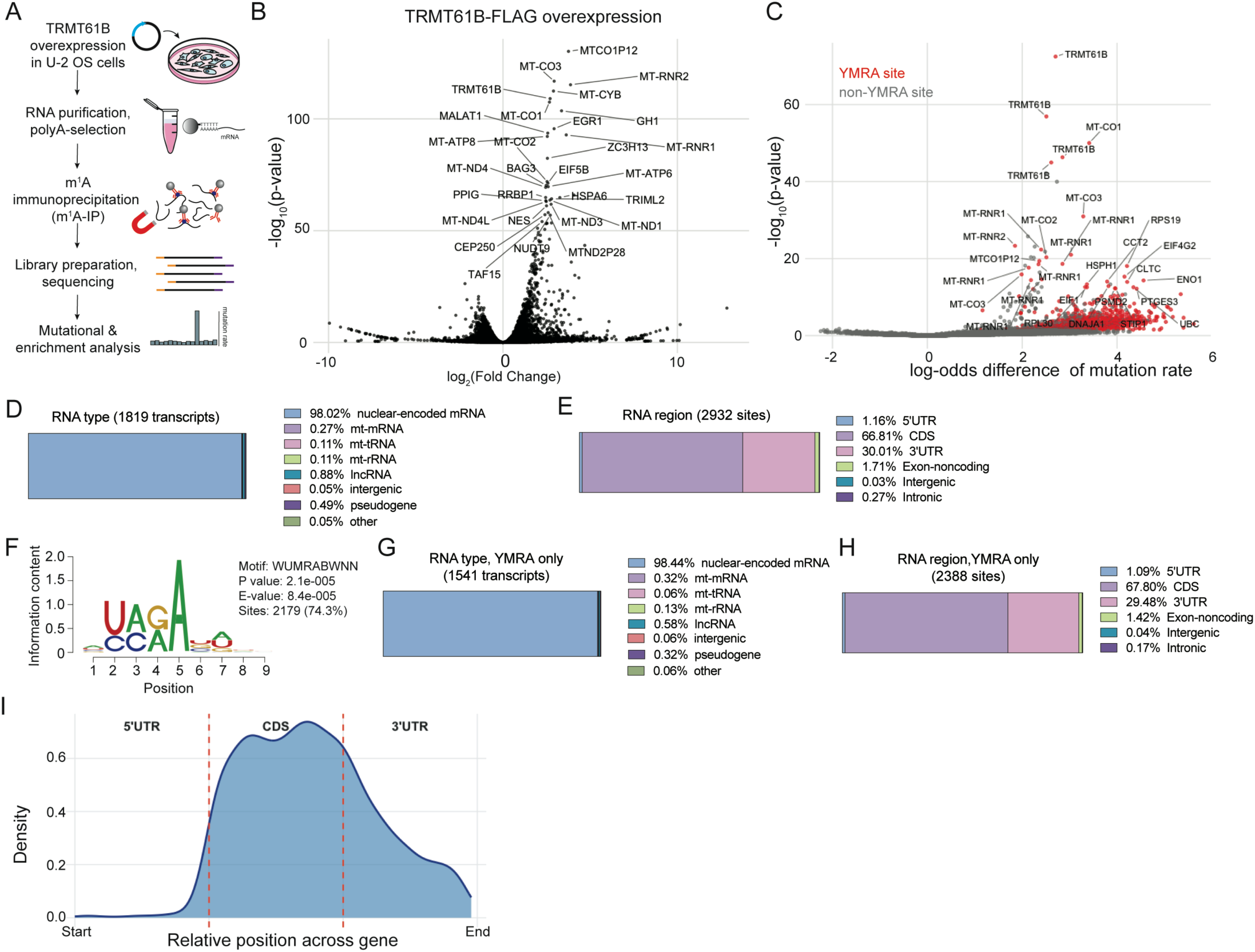
Overexpressed TRMT61B methylates thousands of sites across the transcriptome. A) Schematic of TRMT61B overexpression (OE) in U-2 OS cells and m1A-immunoprecipitation. RNA was purified, polyA-selected, immunoprecipitated, and then prepared into libraries with a TruSeq Stranded mRNA Library Prep kit (Illumina). Both control and TRMT61B overexpression and m^1^A-IP were carried out in duplicate. B) Volcano plot showing DESeq2 [56] results depicting fold changes in transcript abundance in m^1^A-IP samples compared to input samples under TRMT61B overexpression conditions. Top 30 hits by -log_10_(p_adj_) are labeled. C) Volcano plot showing single nucleotide sites called by bakR as having altered misincorporation levels in TRMT61B OE conditions vs control OE conditions, with “high-confidence” YMRA sites (bakR hits with additional filtering of p<0.05, difference in log-odds of mutation rate >1, >1% mutation rate in both replicates, >5% average mutation rate in IP, and <5% average mutation rate in input samples) highlighted in red. Top 30 hits are labeled. D) RNA types of all high confidence single-nucleotide sites, regardless of methylated motif. E) Regions of modification for all high confidence sites, regardless of methylated motif. F) Top motif discovered in high-confidence sites using STREME [55], covering 74.3% of all sites. G) RNA type of high-confidence sites, filtered to only YMRA-motif sites. H) Regions of modification for high-confidence sites, filtered to only YMRA-motif sites. I) Metagene plot showing distribution of YMRA-containing high-confidence sites.

We first analyzed differences at the transcript level between input and m^1^A-IP samples for control and TRMT61B-FLAG overexpressions. As expected, ten of the top 20 most statistically significantly enriched transcripts upon TRMT61B-FLAG overexpression were mitochondrial, including *MT-CO1, MT-CO3* and *MT-RNR2* (16*S* rRNA) (Fig. 1B, Supp. Table 1) [3,5,9]. These transcripts validated our overexpression and immunoprecipitation and confirmed previously reported activity on mitochondrial RNAs. Beyond these mitochondrial targets, many additional RNAs were uniquely enriched in the m^1^A-IP upon TRMT61B overexpression, including *TRMT61B* itself (Fig. 1B, Supp. Fig. 1C). In total, 1,530 transcripts were significantly enriched (p_adj_ <0.05) in the m^1^A-IP relative to input upon TRMT61B-FLAG overexpression (Fig. 1B, Supp. Table 2). Of these 1,530 transcripts, 714 were either not detected or were at least two-fold less enriched in the control m^1^A-IP compared to the TRMT61B-FLAG IP (Supp. Fig. 1C, Supp. Table 2).

To further analyze our m^1^A IP data, we took advantage of the fact that 1) we did not fragment the RNA prior to m^1^A-IP and 2) we prepared sequencing libraries with SuperScript II reverse transcriptase. The positive charge and structure of m^1^A make it disruptive to reverse transcription, and Superscript II has been shown to truncate upon encountering a m^1^A site or to read through it with introduction of a mutation [4,26]. Given that truncations can be difficult to quantify, we chose to complement our transcript-level analysis with mutational analysis on the m^1^A-IP sequencing data. To more robustly analyze mutation signatures while also accounting for potential inter-replicate variability [27], we used bakR, a mutation-calling pipeline that utilizes Bayesian hierarchical modeling to increase statistical power in experiments with low replicate number and variable coverage [28,29]. We identified sites where mutation rates in m^1^A-IP samples were increased compared to input samples under TRMT61B-FLAG overexpression conditions. We then generated a stringently filtered list of high-confidence sites (see methods for criteria) for further analysis. In this high-confidence hit list, thirteen of the top 20 hits were again mitochondrial, including *MT-CO1, MT-CO3,* and *MT-RNR2*, confirming increased m^1^A methylation on known TRMT61B targets (Fig. 1C, Supp. Table 3). We also observed that five of the top 10 hits were sites in the *TRMT61B* mRNA itself. In total, 2,932 high-confidence m^1^A sites were identified across 1,819 different RNAs, including a large number (1,783) of nuclear-encoded mRNAs (Fig. 1D). The majority (64%) of transcripts with m^1^A sites called by mutation signatures carried a single high confidence m^1^A site (Supp. Fig. 1D).

Of the high confidence m^1^A sites in mRNA transcripts, most were located in the coding sequence (CDS) and 3′ untranslated region (UTR) (1,959 and 880, respectively), with only 34 in 5’UTR (Fig. 1E). Motif analysis of sequences surrounding high confidence m^1^A sites revealed a YMR<u>A</u> sequence motif (Fig. 1F; <u>A</u> indicates methylated A). This is consistent with the methylated sites in the known TRMT61B substrates 16*S* rRNA (UAA<u>A</u>) and mitochondrial tRNAs (UCA<u>A</u> / UCG<u>A</u> / CCA<u>A</u>) (Supp. Table 1). Close examination of previously reported mitochondrial mRNA targets of TRMT61B reveals that eight out of nine are also methylated in this motif (Supp. Table 1) [3,5]. Further analysis of only m^1^A sites with this YMRA motif (Fig. 1C, Supp. Table 3) showed that 2,388 out of 2,932 sites were methylated within this motif and that almost all (>98%) of these sites were in nuclear encoded mRNAs (Fig. 1G). These sites were predominantly located in the CDS (67.80%; Fig. 1H, I). CDS m^1^A sites were relatively evenly distributed across codon positions, with a slight preference for position 2 (Supp. Fig. 1E). All codons that could be accommodated within a YMRA motif were represented, but certain codons were underrepresented (serine, methionine) while others were overrepresented (lysine, aspartic acid, glutamic acid, arginine) (Supp. Fig. 1F). Taken together, overexpression of TRMT61B in a human cell line resulted in m^1^A methylation of nuclear-encoded mRNAs, primarily in CDS and 3′UTR regions with a consistent YMRA motif.

### TRMT61B methylates synthetic human RNAs with similar motif preferences

To validate TRMT61B target specificity and discover the downstream effects of mRNA m^1^A methylation on translation, we turned to an orthogonal, *in vitro* system. Our m^1^A-IP data (Fig. 1) revealed TRMT61B-induced m^1^A methylation sites in mRNA CDS regions, and previous published work has suggested that m^1^A presence in CDS regions can reduce translational efficiency [3,30,31]. Additionally, while we observed only a few 5′UTR m^1^A sites in our m^1^A-IP data, we cannot exclude influence arising from the 3′ bias of our sequencing methods (Supp. Fig. 1G) [32]. Several previous studies of m^1^A in human mRNA also found m^1^A to be primarily located in 5′UTR and CDS regions [4,6,20,33]. To investigate the distribution and sequence preferences of TRMT61B m^1^A methylation in more detail and to measure potential downstream effects on ribosome recruitment and translation, we used a diverse pool of *in vitro* transcribed RNAs. This pool contained over 35,000 full-length 5′UTRs and their corresponding early endogenous coding sequences (up to 300 nucleotides total in length) from the human transcriptome, as well as 450 viral 5′UTR/CDS sequences. This pool was previously designed and used to study translation initiation by measuring differential ribosome recruitment [34].

We expressed and purified TRMT61B-FLAG from human cultured cells, and validated its enzymatic activity by methylating a synthetic RNA oligonucleotide containing the sequence of the known m^1^A site in 16*S* rRNA using deuterated *S*-adenosylmethionine (d_3_SAM) as a substrate (Fig. 2A,B). We then measured the resulting levels of deuterated m^1^A (d_3_m^1^A) by LC-MS/MS (Fig. 2A,C). Having confirmed methyltransferase activity on a known substrate, we then methylated our 5′UTR/CDS pool with TRMT61B-FLAG and d_3_SAM under three conditions: no enzyme (control), low enzyme concentration, and high enzyme concentration. We observed dose-responsive TRMT61B-mediated d_3_m^1^A methylation (Fig. 2D), indicating that a subset of the RNA sequences in this pool can be methylated by TRMT61B.

**Figure 2.**
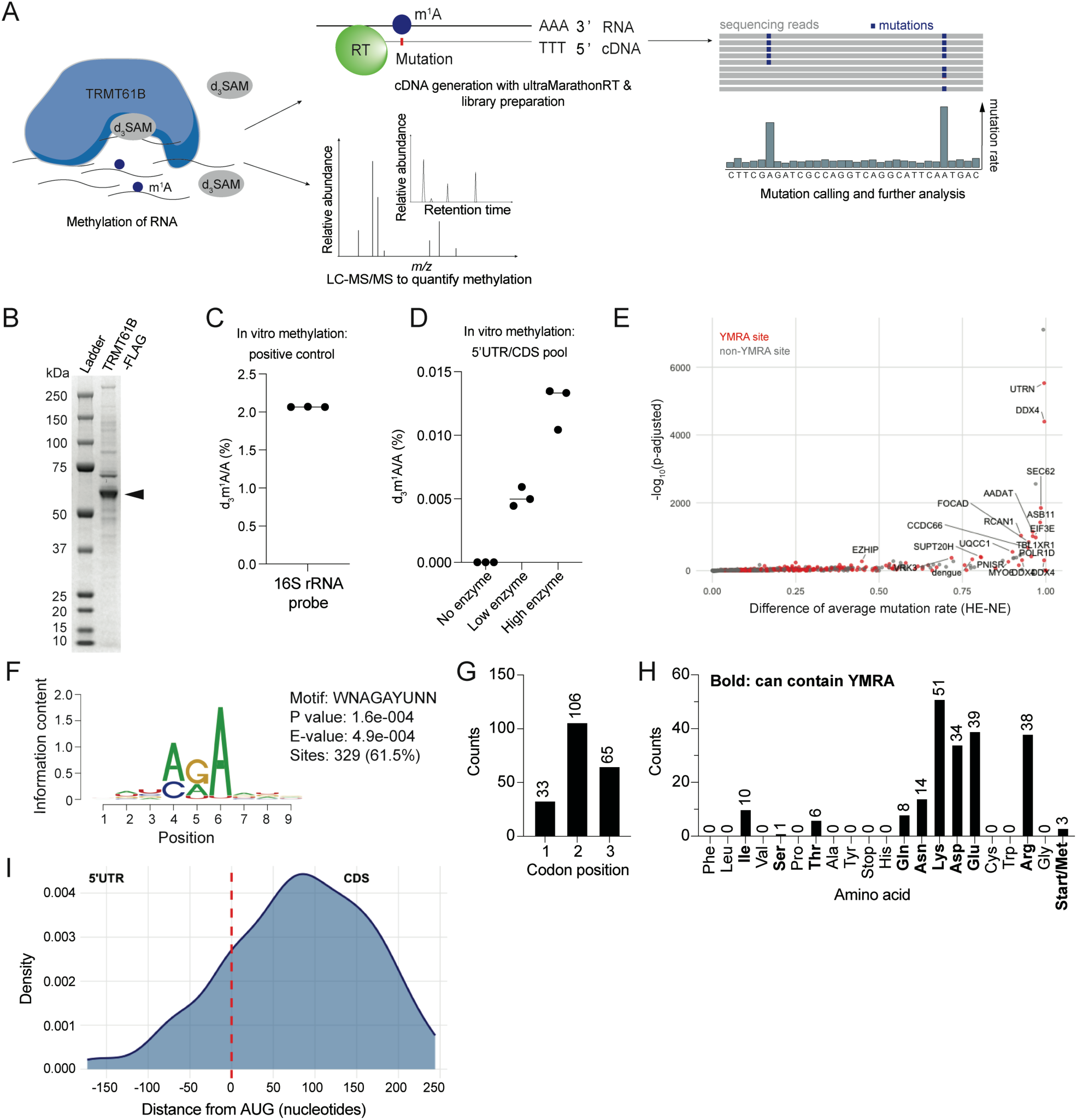
Purified TRMT61B methylates hundreds of sites in a synthetic human mRNA 5’UTR/CDS pool. A) Schematic of *in vitro* TRMT61B methylation of RNA, with validation by LC-MS/MS and detection of single nucleotide sites using ultraMarathonRT-based misincorporation. B) SDS-PAGE gel showing purified TRMT61B protein, indicated by the arrow (∼57 kDa). C) LC-MS/MS measurements of d_3_m^1^A in a short oligo based on 16*S* rRNA (39 nucleotides, centered around adenosine) methylated by purified TRMT61B. D) LC-MS/MS measurements of d_3_m^1^A in a 5’UTR/CDS pool methylated by purified TRMT61B. E) Single nucleotide sites detected in both high and low enzyme 5’UTR/CDS pool samples. “High-confidence” YMRA-containing sites (1% mutation rate in two of three replicates in both HE and LE treatments, padj <0.05 in HE sample) are highlighted in red, with top 20 hits labeled. F) Top motif surrounding high-confidence sites (with duplicated sequences from alternative isoforms removed) using STREME [55], covering 61.5% of all sites. G) Codon positions of high-confidence YMRA single nucleotide sites located in the CDS. H) Distribution of the number of high-confidence 5’UTR/CDS pool YMRA m^1^A sites found in codons of all possible amino acids. Amino acids with codons that can accommodate a YMRA motif are bolded. I) Metagene plot showing distribution of YMRA-containing high-confidence sites across 5’UTR and CDS regions.

To identify the specific sequences from the 5′UTR/CDS pool that were methylated, we used reverse transcription-based mapping of m^1^A sites. To improve m^1^A read through relative to our m^1^A-IP experiments (Fig. 1), we reverse transcribed our *in vitro* methylated 5′UTR/CDS pools using the highly processive ultraMarathonRT (uMRT) enzyme [26]. We sequenced the resulting cDNA at high depth and called m^1^A sites based on increased mutation rates with enzyme treatment in the 5′UTR/CDS pool, resulting in a list of 666 high confidence single nucleotide m^1^A sites (see methods for criteria; Fig. 2E, Supp. Table 4). This list of sequences was de-duplicated to account for alternative isoforms sharing the TRMT61B-methylated sequence, and the resulting 535 unique sequences were used for motif analysis. Motif analysis yielded an AG<u>A</u> motif, similar but less stringent than the motif identified in the TRMT61B-FLAG overexpression experiment (Fig. 2F, compared to Fig. 1F). 61.5% of the m^1^A sites identified in the 5′UTR/CDS pool were found in this motif. Focusing on the sites found in the more stringent YMR<u>A</u> motif, the codon position and amino acid distributions were similar to the m^1^A-IP data, though the bias towards the second codon position was more pronounced in the pool (Fig. 2G,H compared to Supp. Fig. 1E,F). Despite the enrichment of 5′UTR sequence in this pool by design, TRMT61B-mediated methylation was still more prevalent in CDS regions (Fig. 2I), though a higher percentage of 5′UTR sites did appear relative to the m^1^A-IP analysis. Thus, the lower number of 5′UTR sites in our m^1^A-IP (Figure 1) is likely not explained solely by 3′ bias in the sequencing, but faithfully reflects TRMT61B activity on these sequences.

### TRMT61B prefers to methylate the YMRA motif in single stranded regions

The strong agreement in the consensus sequence between the m^1^A-IP and *in vitro* methylated 5′UTR/CDS pool led us to further probe TRMT61B selectivity. We chose seven highly modified transcripts from the *in vitro* methylated pool, choosing sites that were modified under both low- and high-enzyme conditions and that varied in terms of location (5′UTR, CDS) and sequence context (Fig. 3A). Two randomized oligonucleotides were designed per sequence, with the three nucleotides upstream or downstream from the m^1^A site randomized to A/C/G/U in equal proportions (Fig. 3B). These randomized oligos were then *in vitro* methylated with TRMT61B-FLAG and methylation was validated by LC-MS/MS (Supp. Fig. 2A).

**Figure 3.**
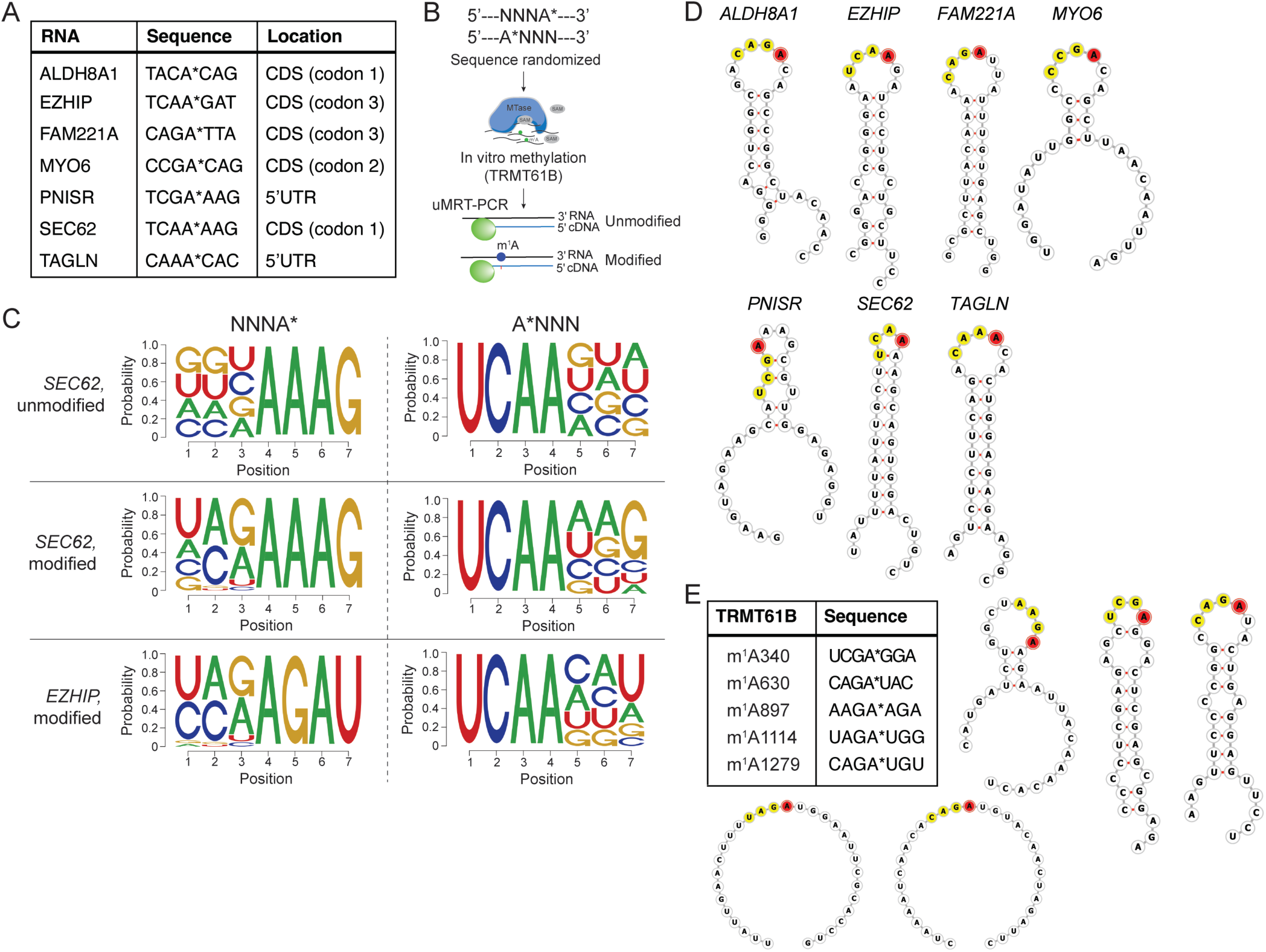
Randomization of methylated sequences confirms upstream YMR sequence preference for TRMT61B. A) Table of the seven mRNA examples chosen for randomized motif experiments, including context sequence and location of single nucleotide site. B) Schematic of randomized motif experiments, showing randomization of upstream and downstream 3-mer followed by methylation and targeted ultraMarathonRT-PCR. C) Example data from *SEC62* and *EZHIP* sequences showing nucleotide abundances at each position between -3 and +3 surrounding the modification site of interest. Amplicon sequencing results were split into “unmodified” samples that had no misincorporation at the modification site, while “modified” sequences contained a misincorporation. Results for the other five transcripts are shown in Supp. Fig. 2. D) Minimum free-energy (MFE) secondary structures surrounding seven example m^1^A sites, generated using RNAfold [35]. The upstream YMR motif is highlighted in yellow, and m^1^A site in red. E) Sequences and MFE secondary structures of five high confidence m^1^A sites identified in *TRMT61B* mRNA under conditions of TRMT61B overexpression.

We then carried out targeted uMRT-PCR and deep amplicon sequencing (Fig. 3B, Supp. Fig. 2B). After checking site-specific methylation, we divided reads into two groups: those with a mutation at the m^1^A site (indicating methylation) and those without a mutation (indicating no methylation). We then analyzed sequence composition of each of these subsets of reads, noting deviations from the initial equal distribution of A/C/G/U (Fig. 3C, Supp. Fig. 2C). Broadly, these results supported the sequence preferences revealed by the consensus motif identified in both the *in vitro* methylation and m^1^A-IP experiments (Fig. 3C, 1F, 2F), including an enrichment in methylated sequences of the YMR motif upstream of the modification site. Interestingly, we also observed strong sequence-dependent selectivity at the +3 position (Fig. 3C) which was not reflected in our earlier experiments. We also quantified 3-mer combinations upstream and downstream of the modification site (Supp. Fig. 2D) and noted that upstream 3-mer combinations were shared across most of our analyzed sequences and were enriched in those fulfilling the YMR combination, but also that there were some exceptions. For example, *FAM221A* showed preferences for GAG or GCG 3-mers and a lack of TCG, CAA, and TAA enrichment, despite the fact that these latter sequences satisfy the motif. Downstream 3-mer combinations showed more flexibility, with gene-specific enriched motifs.

Some RNA modifying enzymes with tRNA targets have structural preferences in addition to sequence constraints, so we also calculated minimum free energy structures for each of these seven sequences using RNAFold [35]. In all cases, we observed that TRMT61B-modified m^1^A sites occur in predicted hairpin loops (Fig. 3D), suggesting a preference for single stranded regions within folded RNAs. Interestingly, though they were not represented in the 5′UTR/CDS pool, the highly m^1^A methylated sites in *TRMT61B* itself that were detected in the m^1^A-IP data also occurred in unpaired RNA regions (Fig. 3E). We also calculated the base pairing probability of all YMRA sites present across the entire synthetic 5′UTR/CDS RNA pool using RNAstructure [36]. We then graphed the distributions of base pairing probabilities at each position in the motif, comparing our high-confidence methylated sites with all other sites. High confidence modified m^1^A sites had a lower probability of base pairing at all positions in the motif, especially at the A of interest (Supp. Fig. 2E). Taken together, these results suggest that the combination of the YMRA sequence and structural preferences could guide TRMT61B to methylate a variety of RNA targets in cells.

### Repression of translation by m^1^A methylation varies by sequence and location

To assess the potential regulatory functions of m^1^A methylation, we incorporated several of the same high confidence, methylated sequences (Fig. 3A) into NanoLuc (Nluc) luciferase reporter constructs (Fig. 4A, Supp. Table 7). In parallel, we generated reporter constructs in which the methylated A was mutated to G, so it could no longer be methylated by TRMT61B-FLAG. This ensured that we could specifically measure the effect of m^1^A methylation at that site, as incubation of these longer reporter RNAs with TRMT61B-FLAG sometimes results in methylation at other sites (Supp. Table 5). Previous studies have suggested that m^1^A in CDS regions has codon position-dependent effects in eukaryotic translation [31]. Given the location of these sites in 5’UTR and early CDS regions, we hypothesized that m^1^A could have differential effects on ribosome recruitment and protein translation.

**Figure 4.**
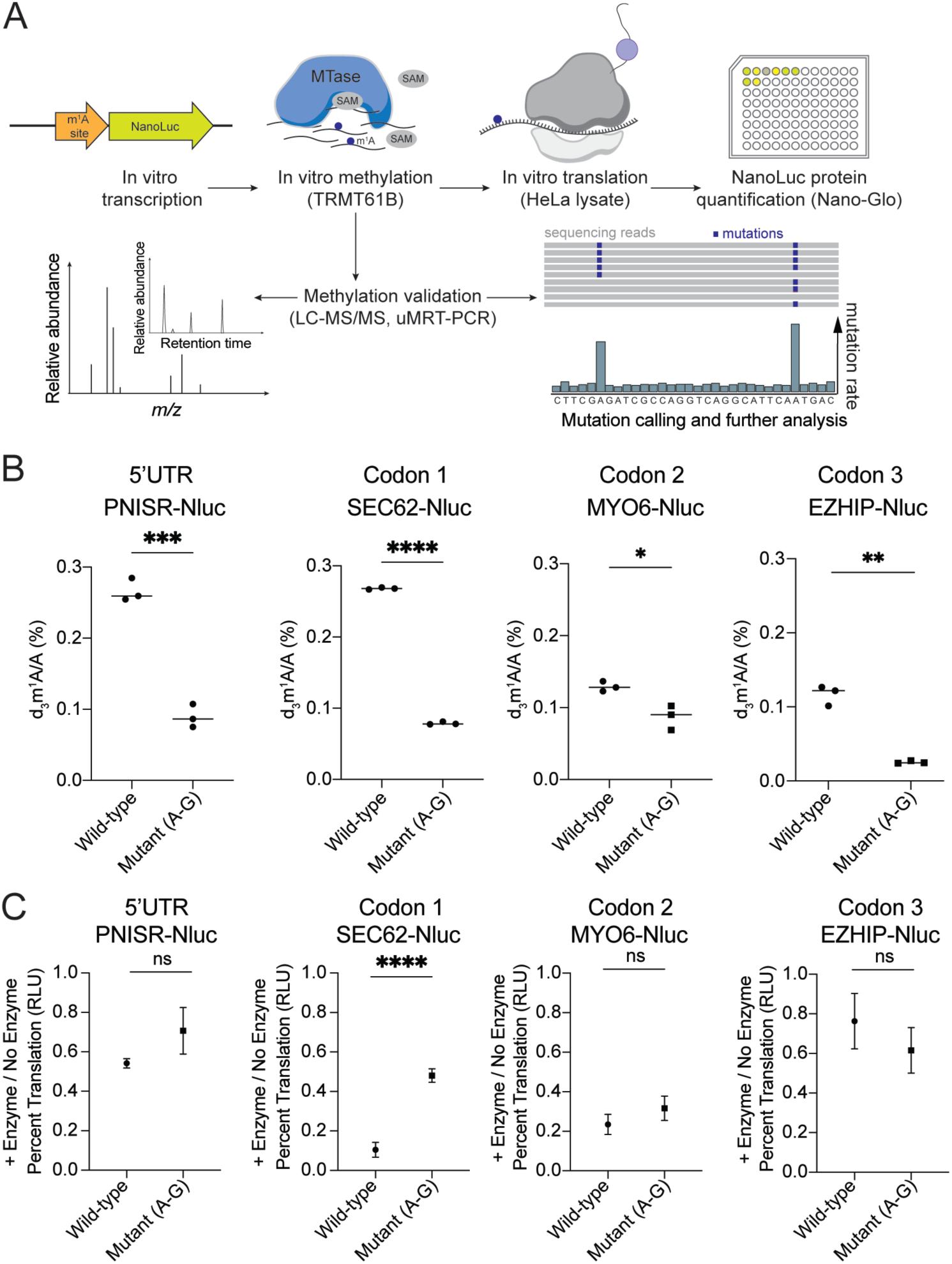
m^1^A modification negatively impacts translation in a context-specific manner, depending on 5’UTR vs. CDS location, codon position, and modification stoichiometry. A) Schematic of experiments measuring *in vitro* translation of *in vitro* transcribed and methylated luciferase reporter sequences. B) LC-MS/MS measurements of d_3_m^1^A levels on wild-type or mutant (m^1^A site A-G mutation) reporters after incubation with purified TRMT61B. Statistical analysis by unpaired t-test, * p<0.05, ** p <0.01, *** p <0.001, and **** p <0,0001 (n=3 technical replicates). C) Luciferase reporter assay results from *in vitro* translation of TRMT61B-treated wild-type or mutant NanoLuc reporters, normalized to translation of untreated wild-type or mutant reporter RNA, respectively. Statistical analysis by two-sample z-test comparing independent ratio estimates using propagated standard errors. * p<0.05, ** p <0.01, *** p <0.001, and **** p <0,0001 (n=3 replicate methylation reactions).

After *in vitro* transcription and methylation of wild-type and mutant luciferase reporter constructs, we validated methylation status by LC-MS/MS (Fig. 4B) and uMRT-PCR (Supp. Fig. 3A). Based on LC-MS/MS measurements, all of the wild-type reporters showed more efficient methylation by TRMT61B-FLAG than their mutant counterparts (Fig. 4B), though each reporter varied in its levels of background d_3_m^1^A, (the result of background methylation in other parts of these longer reporter constructs, example Nluc methylation in Supp. Table 5). Despite this, the effect on *in vitro* translation of our reporters differed significantly. For instance, the PNISR-NLuc (5′UTR site), SEC62-NLuc (CDS, codon position 1), and ALDH8A1 (CDS, codon position 1) reporters had similar bulk levels of d_3_m^1^A (Fig. 4B, Supp. Fig. 3B). While there was no statistically significant difference in the translation of the PNISR-NLuc reporter, methylation reduced translation of the SEC62-NLuc and ALDH8A1-NLuc reporters with CDS methylation sites in a Lys <u>A</u>AA codon and a Thr <u>A</u>CA codon, respectively (Fig. 4C, Supp. Fig. 3B). This effect was less pronounced in the MYO6-NLuc reporter with methylation in an Asp G<u>A</u>C codon, and there was no significant effect with m^1^A in a Gln CA<u>A</u> codon. While the individual reporters are not directly comparable due to their different sequences and methylation levels, these results are consistent with a previous study showing strong to moderate decreases in translation efficiency in a eukaryotic extract by m^1^A methylation in codon positions 1 and 2, but little or no effect in codon position 3 [31].

### m^1^A methylation can alter ribosome recruitment

Our luciferase reporter results are consistent with the idea that m^1^A can have different effects on translation depending on the exact position of the methylation site, particularly when comparing 5′UTR and CDS regions. However, our luciferase assays only effectively read out translation elongation. We hypothesized that while m^1^A in the CDS can be detrimental to translation elongation as has been suggested previously [30,31], m^1^A in the 5′UTR could also potentially impact translation initiation. To test this directly, we used our m^1^A methylated 5′UTR/CDS pools to perform Direct Analysis of Ribosome Targeting (DART) [34]. Briefly, this assay quantitatively measures the recruitment of ribosomes to 5′UTR and early CDS sequences in high throughput, allowing us to compare ribosome recruitment between m^1^A methylated and unmethylated samples. We compared ribosome recruitment in our pools methylated under both ‘low enzyme’ and ‘high enzyme’ conditions (Fig. 2) to unmethylated pools and focused on our set of high confidence YMRA-containing m^1^A methylated sequences (Fig. 2E, Supp. Table 4). The effects of methylation on ribosome recruitment were measured by calculating enzyme treatment-dependent changes in enrichment of transcripts in the 80S ribosome-bound fractions compared to inputs.

From our list of high confidence, m^1^A-methylated YMRA sites in the pool, we detected 73 transcripts with 5′UTR m^1^A sites and 199 transcripts with CDS m^1^A sites in both the input and 80S-bound fractions (Fig. 5A, Supp. Table 6). Analysis of these transcripts showed that for 5′UTR sites, 6 transcripts had significantly increased (p_adj_ < 0.05, log_2_FC > 1) and 28 had significantly decreased (p_adj_ < 0.05, log_2_FC < -1) ribosome recruitment with low or high enzyme treatments, including 5 and 12 unique genes respectively (Fig. 5A). The remaining transcripts were alternative isoforms of the same genes that each contained the consensus YMRA site. For CDS sites, we identified 10 transcripts with significantly increased and 63 with significantly decreased ribosome recruitment in the low and high treatments, with 10 and 33 respective unique genes (Fig. 5A).

**Figure 5.**
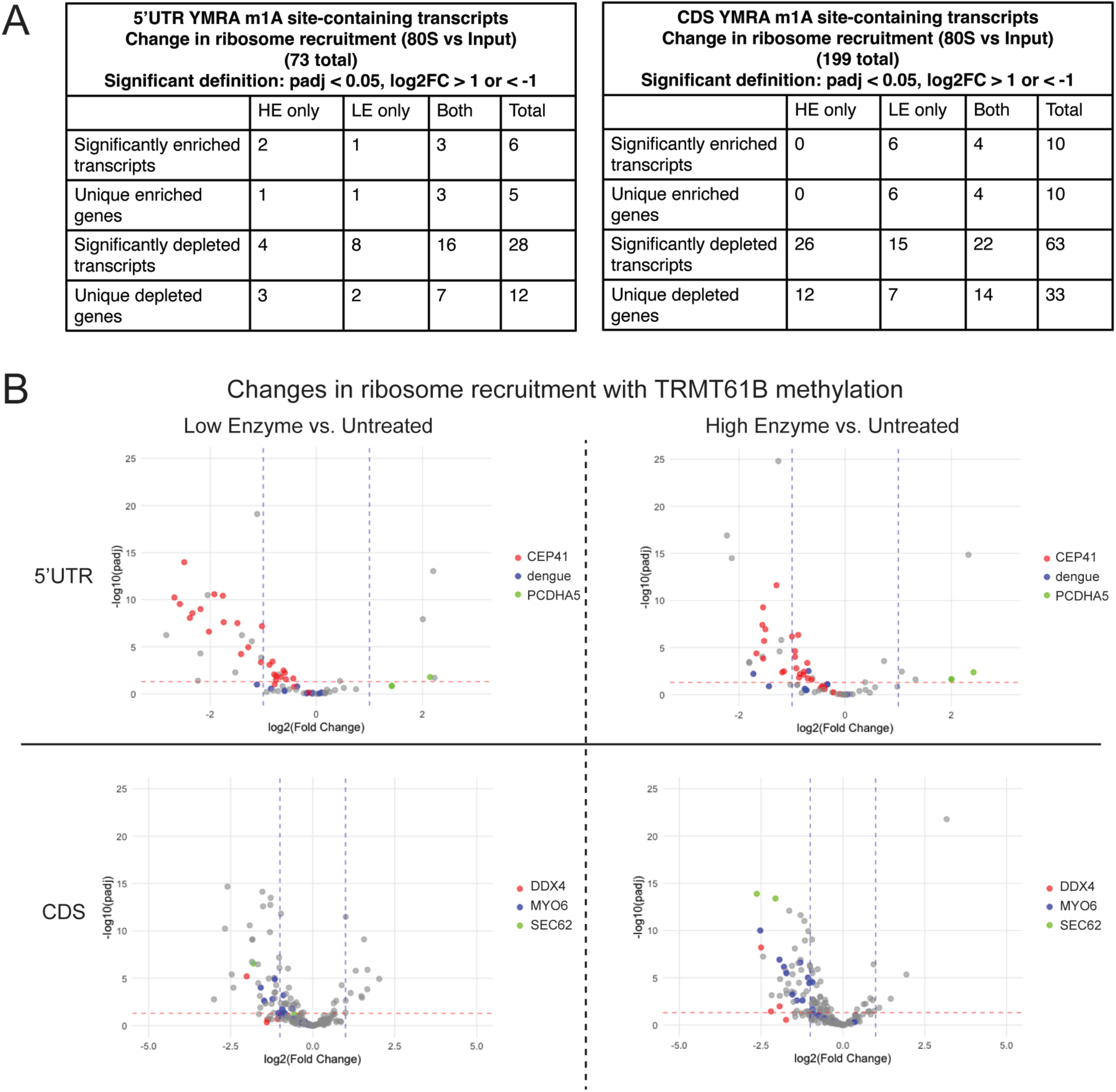
m^1^A modification impacts ribosome recruitment of a synthetic 5’UTR/CDS pool in a transcript-specific manner. A) Table showing counts of transcripts containing high confidence YMRA m^1^A sites with significantly changing ribosome recruitment upon m^1^A methylation (altered enrichment in the 80*S* fraction between TRMT61B- and no enzyme-treatments). Counts are shown for all individual transcripts and by unique genes (which may have multiple enriched/depleted transcript variants). LE = low enzyme, HE = high enzyme. B) Graphs showing log_2_FC in 80*S* enrichment between enzyme-treated and untreated pool samples, specifically depicting transcripts in the pool with high confidence YMRA m^1^A modification sites. Separate graphs are shown for 5’UTR and CDS sites. In each case, three example genes are colored to highlight the consistency in ribosome recruitment changes between transcript variants and between high and low enzyme treatment.

While the overall trends skewed towards m^1^A methylation reducing ribosome recruitment, we observed examples of changes in both directions, suggesting that the effects of m^1^A on ribosome recruitment were transcript-specific. There was strong overlap between significantly altered transcripts in low and high enzyme treatments (Fig. 5B), and also in trends for transcript isoforms of the same gene which all contained the same modified m^1^A site (Fig. 5B). For example, we were able to detect 27 different alternative isoforms of the CEP41 gene that all contained the same m^1^A site in our ribosome recruitment data, and all 27 had decreased ribosome recruitment (negative log_2_FC) with both low and high enzyme treatment (Fig. 5B, Supp. Table 6).

Taken together, these results suggest that a disruptive modification like m^1^A can have highly variable effects on the ribosome recruitment and translation of an mRNA, depending on the specific sequence and structure context. This further highlights the need for detailed, transcript-specific studies of this modification.

## DISCUSSION

In this study, we demonstrate that TRMT61B can methylate hundreds to thousands of non-mitochondrial transcripts, both in a human cell line overexpressing TRMT61B and an *in vitro* methylated synthetic RNA pool (Fig. 1, 2). From the identified single nucleotide sites, we discovered a preferentially methylated YMR<u>A</u> sequence motif in single-stranded RNA regions, and further confirmed the motif through randomization of target sequences surrounding several example m^1^A sites (Fig. 3). We then tested the effects of m^1^A on translation using methylated luciferase reporter constructs and on ribosome recruitment using DART experiments on methylated pools of human 5′UTR and CDS sequences (Fig. 4). We found that m^1^A could significantly impact protein translation, often inhibiting translation and ribosome recruitment. Nevertheless, closer examination of the data on a transcript-by-transcript basis reveals many exceptions, as we also identified sequences where m^1^A methylation did not alter translation or enhanced ribosome recruitment. Taken together, our results highlight the need for detailed characterization of the molecular mechanisms of mRNA modifications on a transcript-specific basis.

Our newly discovered YMRA motif is similar but less stringent than the GUUCRA motif previously discovered for the cytoplasmic tRNA methyltransferase complex TRMT6/61A. Combined with the observation that TRMT61B overexpression can result in activity outside of mitochondria, our findings suggest that TRMT61B may modify nuclear encoded mRNAs in addition to its known mitochondrial targets. Several studies have previously observed primarily mitochondrial localization for TRMT61B [8,37], but that does not rule out its presence outside the mitochondria. Other predominantly mitochondrial proteins have been found to “moonlight” in the nucleus or cytoplasm [38,39], and protein mislocalization is frequently reported in many cancers [40]. Indeed, it remains to be addressed in the future whether such altered expression and localization of TRMT61B occurs, and whether this can induce aberrant m^1^A methylation of non-mitochondrial RNAs. In any case, the TRMT61B sequence preferences and other parameters revealed in our work offer valuable information for facilitating the identification of these potential modification sites. The increased stoichiometry of canonical m^1^A sites in the mitochondria with TRMT61B overexpression that we observe is also consistent with previous work [5] and suggests that aberrant TRMT61B expression in disease could induce changes in mitochondrial function.

Here, we found that TRMT61B preferentially methylated CDS and some 3’UTR sequences, even when presented with a pool that was enriched for 5’UTR sequences (Fig. 2I) [41,42]. Previous antibody-based mapping studies found m^1^A in human mRNA to be primarily localized in the 5’UTR (37-74%) and CDS (19-54%), with the smallest percentage in the 3’UTR (only 1.3-15%) [3,4,6,33]. Later work suggested that the antibody used in those studies had cross-reactivity with the m^7^G-cap [14]. The extent to which this cross-reactivity impacted each study is debated [12,13], so these studies should be interpreted with this caveat in mind. As other work has noted, however, m^1^A deposition is likely cell type- and enzyme-specific [33]. Given likely differences in the evolutionary background of TRMT61B from the other m^1^A methyltransferases [8], it is also not surprising that the distribution of target sites might differ between human methyltransferases. Mapping of m^1^A in other organisms has found varied distributions of this modification, with dinoflagellate m^1^A sites found to be primarily in the 3’UTR and CDS, while petunia m^1^A sites were found primarily in the CDS near the start codon [43,44].

The positional preferences for m^1^A deposition may have implications for the effects of TRMT61B overexpression on translation of downstream target RNAs. Our *in vitro* experiments testing the effects of m^1^A on translation of luciferase reporters and ribosome recruitment in HeLa lysate suggested that m^1^A more often has a negative effect on translation, but that different modified regions of mRNA and exact transcripts vary in their downstream functional consequences. Our tested 5′UTR modification did not result in significantly diminished translation with the addition of m^1^A, even when the level of modification was similar to a CDS site in the first codon position that caused a 2 to 4-fold drop in translation relative to a mutant control. However, another tested CDS site in the third codon position did not affect translation. These results are consistent with previous reports that also observed inhibitory effects of m^1^A on translation in a variety of organisms (yeast, human cells, bacteria, dinoflagellate, plants), but differing results dependent on RNA region and codon position [3,5,44–46]. A recent study testing a targeted mitochondrial demethylase (MTS-PUF-ALKBH3 fusion protein) found that in two cell lines, removal of the m^1^A in *MT-ND5* increased protein levels, but that the opposite occurred for *MT-CO1* [47]. These results support our transcript-specific findings, and even suggest that for certain transcripts m^1^A can increase translation.

We further explored these context-dependent effects on translation with experiments using a TRMT61B-treated synthetic 5′UTR/CDS pool to measure ribosome recruitment with and without m^1^A. We found that ribosome recruitment could significantly increase or decrease with the addition of m^1^A (with a slight bias towards decreasing), depending on the transcript of interest. Trends for individual transcripts were generally consistent between low and high enzyme treated samples and between alternative isoforms of a single gene, supporting the validity and replicability of our findings. This is the first experiment directly measuring ribosome recruitment for translation initiation to m^1^A-methylated transcripts, and our results suggest that m^1^A can affect translation initiation in addition to the effects on elongation observed previously and in our experiments here [30,31].

Our synthetic mRNA pool did not include any 5′UTR sequences, due to its design as a pool to study ribosome recruitment to 3′UTR sequences, so our conclusions cannot extend to the effects of m^1^A in the 5′UTR. Other work has found that 5′UTR and CDS sites but not 3′UTR m^1^A methylation affected mRNA translation [3], and that 3′UTR m^1^A sites in an mRNA can lead to increased RNA decay or altered binding of miRNAs [33,48,49]. Our findings are consistent with a model of m^1^A having highly variable and context-specific effects on mRNA. Future work utilizing more diverse sequences is needed to draw conclusions about the entire length of an mRNA transcript.

Given the particularly disruptive nature of m^1^A to RNA structure and base pairing, it is not surprising that we sometimes observe drastic changes in translation or ribosome recruitment, or that our results would be highly sequence specific. m^1^A has been shown to interfere with both Watson-Crick base pairing and Hoogsteen pairing in A-RNA, which would interfere with RNA folding and with the interactions between mRNAs and tRNAs during decoding [50]. In addition to enzymatically added m^1^A, alkylation damage can also manifest as m^1^A in RNA [51]. These adducts have been shown to cause ribosome stalling and induce of protein and RNA quality control mechanisms, and left unrepaired they can cause DNA double-stranded breaks [46,51]. These findings suggest that m^1^A is a modification that must be strictly regulated, and that further research into m^1^A may be critical to understanding what happens when it becomes misregulated. Despite or even because of its overall low presence in mRNA, its misregulation may dramatically shift the dynamics of modified transcripts.

One surprising finding in our experiments was that when overexpressed, *TRMT61B* was methylated at five different sites in its own mRNA coding sequence. Given potential negative effects of CDS m^1^A methylation on translation, it is possible that elevated levels of the protein may induce autoregulation where TRMT61B methylates its own RNA to decrease expression. Similar feedback has been observed for other modifying enzymes, including those that regulate m^6^A in mRNA. For example, it has been shown that silencing of ALKBH5 (an m^6^A demethylase) led to reduced levels of METTL14, METTL3, and WTAP (m^6^A writer complex components), and that overexpression of METTL14 or ALKBH5 led to reciprocally increased expression of the other protein, suggesting a tightly regulated feedback loop to control m^6^A levels in the cell [52]. Further study of these autoregulatory feedback loops in disease contexts where enzymes become dysregulated could yield further insights into how RNA modifications impact a variety of pathologies.

Taken together, our study presents a detailed characterization of the TRMT61B m^1^A methylation motif and suggests the potential ability of TRMT61B to act as an m^1^A methyltransferase outside of the mitochondria. Our newly discovered motif and structural preferences will help inform future studies of TRMT61B targets, particularly in settings where TRMT61B expression is misregulated. Additionally, we present *in vitro* evidence that m^1^A presence can significantly affect translation, and emphasize that further work towards understanding this regulation needs to be highly transcript-specific until regulatory pathways for m^1^A are better established. More broadly, our work establishes new possibilities for what the TRMT61B enzyme *can* do, laying the groundwork for more detailed and better controlled experiments in cells and organisms.

## METHODS AND MATERIALS

### Cell Culture

U-2 OS cells (HTB-96) were obtained from the American Tissue Culture Collection (ATCC) and maintained in McCoy’s 5A Medium (Gibco, 16600082) with 10% fetal bovine serum and 1X penicillin-streptomycin at 37°C under a humidified atmosphere with 5% CO_2_. FreeStyle™ 293-F Cells (R79007) were obtained from Invitrogen and maintained in FreeStyle™ 293 Expression Medium (Gibco, 12338018), shaking at 37°C under a humidified atmosphere with 8% CO_2_.

### FLAG-tagged TRMT61B expression and purification

293-F cells were transfected with 293fectin (Invitrogen, 12347019) and TRMT61B plasmid (Origene RC201499) following manufacturer instructions. In short, cells were diluted to 1e6/mL in fresh media, for a total volume of 1 L cells. 1000 μg DNA and 2 mL 293fectin were added to Opti-MEM (Gibco, 31985070) media for 5 minutes, mixed together and incubated for 20 minutes, then added to cells. After 64 hours, cells were harvested by centrifugation at 200 x g for 5 minutes, washed once with 1X ice-cold PBS, and resuspended in 5 pellet volumes of ice-cold lysis buffer made with 50 mM Tris-HCl (pH 7.5), 50 mM NaCl, 1.5 mM MgCl_2_, 0.5 mM CaCl_2_, 10% NP-40, 10% glycerol, and 1X SigmaFast Protease Inhibitor (Millipore-Sigma, S8830). Cells were rotated in lysis buffer for 1 hour at 4°C, and then spun down at 14,000 x g for 20 minutes at 4°C. Supernatant was transferred to a new tube and spun again. Clarified supernatant was incubated with pre-washed Anti-FLAG® M2 beads (Millipore-Sigma, M8823), rotating for 1.5 hours at 4°C. Beads were then washed 2X with lysis buffer, 3X high salt buffer (50 mM Tris-HCl pH 7.4, 500 mM KCl, 0.05% NP-40), and 1X with elution buffer (50 mM Tris-HCl pH 7.4, 150 mM KCl, 1.5 mM MgCl_2_, 0.1 mM EDTA, and 1X SigmaFast Protease Inhibitor). Beads were then eluted with 1 mg/mL 3X FLAG peptide (Millipore-Sigma, F4799-25MG) in 1X elution buffer, rotating for 1 hour at 4°C. Supernatant was then transferred to a 30 kDa Amicon ® Ultra Centrifugal Filter (Millipore-Sigma, UFC903024) and centrifuged for 3-5 minutes at a time at 4,000 x g and 4°C. Spins were repeated 7-10X, with addition of fresh elution buffer each time to remove FLAG and buffer exchange. Final protein was stored in 1X elution buffer and 20% glycerol. Purified protein quality was analyzed on a NuPage 4-12% Bis-Tris gel (Invitrogen, NP0335BOX) followed by staining with GelCode™ Blue Coomassie stain (Thermo Scientific, 24590).

### Western Blot

Cells were washed once with cold 1X PBS and then scraped from plates into 1 mL cold PBS and pelleted at 1,000 x g for 8 minutes at 4°C. Supernatant was discarded after centrifugation and cell pellets were lysed on ice for 30 minutes in three pellet volumes of 1X lysis buffer (50 mM Tris pH 7.4, 300 mM NaCl, 1% NP-40, 0.25% deoxycholate, 1 mM DTT, and 1x SigmaFast protease inhibitor). After 30 minutes, lysate was centrifuged at 20,000 x g for 20 minutes at 4°C, and supernatant was collected. Protein concentration was measured using the Pierce™ 660nm Protein Assay Kit (Thermo Scientific, 22660), and then desired amounts of protein were loaded with 1X protein loading dye (45 mM Tris pH 6.8, 2% SDS, 6% glycerol, bromophenol blue, 100 mM DTT) onto 4-12% NuPage Bis-Tris gradient gels (Invitrogen) and run at 200 V for 50 minutes in 1X MOPS buffer. Protein was transferred to a nitrocellulose membrane (Bio-Rad, 1620112) by semidry transfer at 17V for 45 minutes using 2X NuPage transfer buffer (50 mM Bicine, 50 mM Bis-Tris, 2 mM EDTA, pH 7.2) and 10% methanol. The membrane was then blocked for one hour with 5% milk in TBS then rocked overnight at 4°C with primary antibodies diluted in 5% milk in TBST. Anti-GAPDH (R&D Systems, MAB5718) antibody and anti-TRMT61B (Atlas Antibodies, HPA026751) were both used at 1:1000 dilutions. After primary incubation, the membrane was washed 3 x 5 minutes in TBST, and then incubated for two hours at RT in secondary antibodies diluted in 5% milk in TBST and imaged on a ChemiDoc MP Imaging System (Bio-Rad). Donkey anti-Rabbit IgG secondary antibody DyLight 800 and Donkey anti-Mouse IgG secondary antibody DyLight 680 (Invitrogen, SA510044 and SA510170) were both used at 1:5000 dilutions.

### Quantitative real-time PCR

500 ng of total RNA was reverse transcribed with iScript Reverse Transcription Supermix for RT-qPCR (Bio-Rad) according to the manufacturer’s instructions. 1 μL of 5-fold diluted cDNA, 200 nM of forward/reverse primer, and 5 μL SsoAdvanced Universal SYBR Green Supermix (Bio-Rad, 1725274) were combined per reaction, carried out in triplicate per cDNA/primer combination. RT-qPCR was performed using the Bio-Rad CFX96 Real-Time PCR System (Bio-Rad), and 2-ΔΔCT method and using normalization to GAPDH or HPRT1. Primer pairs are listed in Supp. Table 7.

### RNA purification

Cells were washed once with cold 1X PBS and then lysed in TRIzol reagent (Invitrogen, 15596018) for 3-5 minutes. TRIzol lysate was then transferred from dishes to Eppendorf tubes and 20% volume added of chloroform, with the mixture vigorously shaken for 30 seconds, followed by rest at RT for 3 minutes. The TRIzol-chloroform mixture was then centrifuged at 12,000 x g for 15 minutes at 4°C. Aqueous phase was transferred to clean nuclease-free Eppendorf tubes, followed by addition of 1 μL GlycoBlue (Thermo Fisher, AM9516) and 50% of the original lysate volume of isopropanol. After isopropanol precipitation at room temperature for 10 minutes, RNA was pelleted for 15 minutes at 15,000 x g at 4°C. RNA pellets were washed once with 80% ethanol, dried, and resuspended in RNAse-free water. PolyA-RNA selection was then carried out using the NEBNext® High Input Poly(A) mRNA Isolation Module (NEB, E3370S), following manufacturer protocols.

### RNA modification quantification by LC-MS/MS

For analysis by liquid chromatography-tandem mass spectrometry (LC-MS/MS), 50-100 ng of RNA was digested for 3 hours at 37°C with 10 U Benzonase Nuclease (Millipore-Sigma, E8263-5KU), 0.1 U alkaline phosphatase (Millipore-Sigma, P5521-2KU), and 1 U phosphodiesterase I (Millipore-Sigma (P3243-1VL), in buffer containing 50 mM Tris-HCl (pH 6.5) and 2 mM MgCl_2_. Samples were then filtered with Millex 0.22 μm syringe filters (Millipore, SLGVR04NL) by centrifugation at 10,000 x g for 1 minute. Liquid chromatography was performed on a Shim-pack GIST C18 2 μm, 2.1 × 50 mm column (Shimadzu, 227-30001-02), using a binary gradient with 0.1% aqueous formic acid to 50% methanol with 0.1% formic acid over 8 min at 400 μL/min. Mass spectrometric measurements were performed with a Shimadzu Scientific Instruments 8060 Triple-Quad LC-MS system, equipped with a Nexera LC-40D xs UHPLC, consisting of a CBM-40 Lite system controller, a DGU-405 Degasser Unit, two LC-40D XS UHPLC pumps, a SIL-40C XS autosampler and a Column Oven CTO-40S. UV data was collected with a Shimadzu Nexera HPLC/UHPLC Photodiode Array Detector SPD M-40 in the range of 190 - 800nm. Mass spectra were subsequently recorded with the triple quadrupole (TQ) 8060 mass spectrometer. The ionization source was run in ESI mode, with the electrospray needle held at +4.5kV. Nebulizer Gas was at 2 L/min, Heating Gas Flow at 10 L/min and the Interface at 300°C. Dry Gas was at 10 L/min, the Desolvation Line at 120°C and the heating block at 400°C. Nucleosides were identified using specific nucleoside-to-base ion transitions (284-to-152 for G, 268-to-136 for A, 285-153 for d_3_m^1^A and d_3_m^6^A, and 298-to-166 for m^7^G) and retention times (1 min, 0.9 min, 0.6 min, 2.2 min, 0.7 min, respectively) and quantified using standard curves of pure nucleosides. Measurements and data post-processing were performed with LabSolutions 5.97 Realtime Analysis and PostRun, and with LabSolutions Insight Version 3.7.

### TRMT61B overexpression

Myc-DDK-TRMT61B overexpression plasmid and negative control plasmid were obtained from Origene (RC201499, MR227353). The day before transfection, 1.5 million U-2 OS cells were plated in 15 cm^2^ tissue culture dishes. The next day, media was changed to media without antibiotics and cells were transfected using Lipofectamine 3000 (Invitrogen), following manufacturer protocols. In short, 15 μg of plasmid DNA and 45 μL Lipofectamine 3000 were combined with 30 μL P3000 reagent in Opti-MEM media, incubated for 15 minutes, and then added to cells. After 48 hours, cells were harvested for protein and RNA extraction. Transfections were carried out in duplicate.

### m^1^A-IP and library preparation

6 μL Dynabeads Protein G (Invitrogen, 10004D) was washed three times with PBST (PBS + 0.02% Tween-20), then incubated with 1.5 μg of anti-m^1^A antibody (Abcam, ab208196) for 1 hour on ice with periodic mixing. Beads were then washed once with PBST and twice with 1X IP buffer (10 mM Tris-HCl pH 7.4, 50 mM NaCl, 0.1% NP-40). 1.5 μg polyA-selected RNA and 80 U Protector RNAse inhibitor (Roche,3335399001) in 1X IP buffer was then added to the antibody-bead complex in 1X IP buffer and rotated for 1 hour at 4°C. Following incubation, beads were washed six times with 1X IP buffer. RNA was then eluted by shaking beads in 50 μL TRIzol reagent (Invitrogen, 15596018) for 5 minutes twice. TRIzol eluate was purified with Direct-zol Microprep kit (Zymo, R2061), following manufacturer instructions. cDNA library preparation was performed using TruSeq Stranded mRNA Library Prep kit (Illumina, 20020594). Paired-end 100 bp sequencing was carried out on a NovaSeq X Plus instrument (Illumina), with 50 million read pairs per sample.

### Data analysis

Raw fastq files were processed with Trimmomatic [53] for adapter removal and quality trimming, followed by alignment with STAR [54] using the parameters: --alignSJoverhangMin 8 -- outFilterMultimapNmax 20 --alignSJDBoverhangMin 1 --outFilterMismatchNmax 999 -- outFilterMismatchNoverReadLmax 0.04 --alignIntronMin 20 --alignIntronMax 1000000 -- alignMatesGapMax 1000000. Bamfiles were then sorted with samtools and mutation counts tables were generated with fastq2EZbakR [29].

Sites where mutation rates in the IP were significantly increased over input samples were then identified with bakR [28]. For high-confidence sites, additional filtering cutoffs of average <5% mutation in input samples and >5% mutation in IP samples, individual replicates having >1% mutation rate in IP samples, p-value <0.05, and difference in log-odds of mutation rate >1 were used. Motif analysis was performed with STREME [55] on a deduplicated list of statistically significant sites. Exact parameters are as follows: --verbosity 1 --oc. --rna --totallength 2500000 --time 14400 --minw 7 --maxw 9 --thresh 0.05 --align center. For further analysis, significant sites were filtered for only YMRA-motif containing sequences. Analysis of transcript enrichment in all IP treatments versus input treatments were carried out using DESeq2 [56].

### DART library design

The 300 nucleotide length 5’UTR/CDS RNA oligo pool used for *in vitro* methylation was designed as described in Lewis et al. 2025 [34], with only sequences containing endogenous CDS used. In total, there were 35,409 human mRNA sequences and 464 viral mRNA sequences. The designed pool was commercially purchased (Twist Bioscience) as DNA, then PCR amplified with barcoded reverse primers. Four barcoded primers were used to amplify the pool for each treatment (no enzyme, low enzyme, and high enzyme), with one additional barcode used when amplifying the spike-in 16*S* positive control RNA (Supp. Table 8). DNA pools were then vitro transcribed with co-transcriptional capping using CleanCap AG (Trilink, N-7113) using the MEGAshortscript T7 transcription kit (Invitrogen, AM1354).

### DART library methylation

Capped pool RNA was incubated with no enzyme, low, or high concentrations of purified TRMT61B in 1X methyltransferase buffer (10 mM HEPES pH 7.4, 100 mM KCl, 1.5 mM MgCl2, 1 mM DTT), with addition of 40 U Protector RNAse inhibitor (Roche), 1 mM d_3_SAM (CDN Isotopes), and up to 4% glycerol. One large (640-680 μL) reaction was carried out for TRMT61B methylation site identification and as input for six DART translation reaction replicates, and two additional 40 μL reactions were carried out for additional TRMT61B methylation site identification replicates. “No enzyme” control reactions were carried out with only methyltransferase storage buffer (see purification methods). Following methylation, RNA was purified by gel extraction from a 6% TBE-Urea gel (Invitrogen). Gel slices were excised and incubated for 4 hours at RT in 200 μL gel elution buffer (20 mM Tris pH 7.5, 1 mM EDTA, 250 mM sodium acetate, 0.25% SDS), including 3X freeze-thaw on dry ice for 15 minutes each, followed by TRIzol extraction and purification as described above.

### Library preparation for mutation analysis

For modification detection, 400 ng of RNA from each treatment was then denatured at 65°C for 5 minutes and then reverse transcribed using ultraMarathonRT in 1X buffer containing 50 mM Tris-HCl (pH 7.5), 200 mM KCl, 2 mM MnCl_2_, 5 mM DTT, and 20% glycerol, with 500 μM of dNTPs and 500 nM of oKA210 primer (Supp. Table 8) added. After reverse transcription for 3 hours, 2 μL freshly prepared 1M NaOH was added and the sample was heated to 98°C for 20 minutes, followed by neutralization with 1M HCl. Samples were then run on an 8% denaturing PAGE gel and gel slices were excised and incubated with DNA elution buffer (300 mM NaCl, 10 mM Tris-HCl pH 8.0). DNA eluate was then purified by isopropanol precipitation, with final pellet resuspension in 5 μL Tris-HCl pH 7.5. 0.8 μL of 100 μM DART_5’Adapter (Supp. Table 8) and 1 μL DMSO were added and samples were heated to 75°C for 2 minutes, followed by ligation with High Concentration T4 RNA Ligase I (NEB, M0437M), shaking overnight at RT. Ligated cDNA was then purified using 10 μL of MyOne Silane beads (Invitrogen, 37002D) and libraries were then amplified in an initial round of PCR using an i5 primer and oKA182 (Supp. Table 8). Excess primers were removed by treating 50 μL PCR reactions with 3 μL of thermolabile exonuclease 1 (NEB, M0568L) for 4 minutes at 37 °C before heat inactivating for 1 min at 80 °C. Amplified DNA was then purified using a Monarch Spin PCR & DNA Cleanup Kit (5μg) (NEB, T1130S). Purified DNA was used as a template for the final library PCR reactions using the necessary i5 and i7 barcoded library primers. Final libraries were concentrated using AMPure XP beads (Beckman Coulter, A63881) before purifying on native 8% polyacrylamide gels.

### Data analysis for mutational calling

One round paired end 150 bp sequencing and two rounds of paired end 250 bp sequencing was carried out on a NovaSeq 6000 and NextSeq 2000 respectively (Illumina) for *in vitro* methylated pool samples for mutation analysis (∼25 million reads/sample for 2 x 150, ∼130 million reads/sample for 2 x 250). Raw fastq files were deconvoluted with Flexbar [57], followed by trimming of adapters, removal of UMIs, and collapsing of duplications with BBduk [58]. Reads were then aligned with STAR with the following parameters: --alignIntronMax 1 --alignIntronMin 1 --alignSJoverhangMin 1 --outFilterMismatchNmax 3 --outFilterMismatchNoverReadLmax 0.1 --alignEndsType Local --outFilterMatchNmin 10 -- seedSearchStartLmax 50 --seedPerReadNmax 10000 --outFilterMatchNminOverLread 0.25 -- outFilterScoreMinOverLread 0.25. Counting of alignment was performed with BBMap pileup [58], followed by bamfile conversion and sorting with samtools [59]. Bamfiles for three rounds of sequencing were merged by sample. Single nucleotide sites were then identified using an ad-hoc cutoff-based pipeline. High confidence sites were then selected as those common to both low and high enzyme treatments, with cutoffs of 1% mutation rate in at least two of three replicates in both treatments and an adjusted p-value of <0.05 in the high enzyme samples. Motif analysis was performed with STREME [55] on a deduplicated list of statistically significant sites, using the full pool sequences as background, Exact parameters are as follows: --verbosity 1 --oc. --rna --totallength 100000 --time 14400 --minw 9 --maxw 9 --thresh 0.05 --order 1 --align center --n m1Agenome_EndogenousPool.fasta. For further analysis, sites were additionally filtered to only those containing a YMRA motif at the modification site.

We also globally identified YMRA motifs (Y = C/T, M = A/C, R = A/G, A) across all sequences in the pool after removing the T7 promoter sequence (GCTAATACGACTCACTATAAGG). For each identified motif, we computed the RNA secondary structure ensemble using RNAstructure [36] and calculated the base-pairing probability of each individual nucleotide in the YMRA motif. We then split out base pairing probabilities for our high confidence m^1^A sites.

### TRMT61B DART and library preparation

*In vitro* translation, sucrose gradient preparation, 80S fraction isolation, RNA purification were carried out as described in Lewis et al. 2025 [34]. In total, four replicate *in vitro* translation reactions were carried out in a volume of 500 μL using 40 pmol of pooled RNA containing equimolar ratios of no, low, and high enzyme-treated RNA. Input pooled RNA and purified 80S-bound RNA were then reverse transcribed using Induro RT (NEB, M0681) using oKA210 primer. Following reverse transcription at 60°C for 1 hour, samples were processed as described above for the mutational analysis libraries. Input libraries were prepared in triplicate from pooled RNA, and four 80S fraction libraries from four DART replicates.

### Data analysis for ribosome recruitment

Paired end 150 bp sequencing was carried out on the NovaSeq 6000 (Illumina) for *in vitro* methylated input and 80S fractions for ribosome recruitment analysis (25 million reads/sample). Raw fastq files were processed and aligned as described above for the mutation analysis libraries. Comparisons of transcripts in input and 80S fractions were carried out using DESeq2 [56], with ribosome recruitment analysis carried out on all transcripts that contained at least 10 reads in the input samples and at least 1 read in the 80S fraction. Interaction terms were calculated based on changes in RNA transcript enrichment in the 80S between the no enzyme and low or high enzyme treatment, allowing for analysis of changes in ribosome recruitment induced by addition of low or high TRMT61B enzyme treatment.

### Targeted *in vitro* oligo design

m^1^A modification sites in seven modified RNAs of interest (*ALDH8A1, EZHIP, FAM221A, MYO6, PNISR, SEC62, TAGLN*) were selected for motif randomization and *in vitro* luciferase translation experiments. For motif randomization experiments, 150-250 nucleotide sequences surrounding the m^1^A sites were commercially purchased (IDT) with common RT-PCR handle sequences and bases either 3 nucleotides upstream or downstream of the modification site were randomized to 25% A/C/G/T (Supp. Table 7). For *in vitro* luciferase translation experiments, 150-250 nucleotide sequences surrounding the m^1^A sites were commercially purchased (IDT, Supp. Table 7) and inserted downstream of a T7 promoter and upstream of a NanoLuc sequence cloned into a pT7CFE1-His plasmid (Invitrogen, 88860) with the native T7 promoter mutated to be nonfunctional (TAATACGAgagACTATA) and cut with NotI-HF (NEB, R3189S). Full 5’UTRs were included for 5’UTR m^1^A modification sites, while coding sequences were truncated to reach a similar length. Both wild-type and mutant plasmids (A-G mutation at modification site) were cloned, and plasmids were linearized with SpeI-HF (NEB, R3133S) prior to *in vitro* transcription.

### *In vitro* transcription

Short RNA oligos for motif randomization experiments (<300 nucleotides) were transcribed from PCR products using the MEGAshortscript™ T7 Transcription Kit (Invitrogen, AM1354). Transcribed RNA was then run on a 6% TBE-Urea gel. Gel slices were excised and incubated overnight at 4°C in gel elution buffer (1 mM EDTA, 300 mM sodium acetate), followed by purification of RNA from the supernatant by isopropanol precipitation. Longer, capped RNA oligos for *in vitro* translation assays were transcribed using the mMESSAGE mMACHINE™ T7 mRNA Kit with CleanCap™ Reagent AG (Invitrogen, A57620). Transcribed RNA was then run on a 6% TBE-Urea gel. Gel slices were excised and incubated between 3-16 hours at RT in 200 μL gel elution buffer (20 mM Tris pH 7.5, 1 mM EDTA, 250 mM sodium acetate, 0.25% SDS), including 3X freeze-thaw on dry ice for 15 minutes each. RNA was then purified by TRIzol extraction, as previously described. All RNA quality was analyzed using a High Sensitivity RNA ScreenTape (Agilent, 5067-5579) run on an Agilent 4200 TapeStation System.

### Targeted *in vitro* methylation reactions

*In vitro* methylation was carried out by combining target RNA and purified TRTM61B protein in 1X methyltransferase buffer (10 mM HEPES pH 7.4, 100 mM KCl, 1.5 mM MgCl_2_, 1 mM DTT) with addition of 40 U Protector RNAse inhibitor (Roche, 3335399001) and 1 mM d_3_SAM (CDN Isotopes, D- 4093). Glycerol in the reaction was adjusted to a final amount of 4%, accounting for glycerol in purified protein added. Reactions were incubated for one hour at 37°C. Final reactions were cleaned up with an RNA Clean and Concentrator kit with size selection for large RNAs (Zymo, R1016), following manufacturer instructions.

### Targeted *in vitro* translation assays

*In vitro* translation reactions were carried out using the 1-Step Human Coupled IVT Kit (Invitrogen, 88882), following manufacturer instructions. In short, 100-250 ng of *in vitro* transcribed, capped RNA was mixed with kit reagents and incubated for 2-4 hours at 30°C. Luciferase activity was then measured using the Nano-Glo Luciferase Assay System (Promega, N1150).

### Targeted modification detection by uMRT-PCR

RNA (100 ng for *in vitro*-transcribed RNAs) was reverse transcribed with ultraMarathonRT (RNAConnect) in 1X buffer containing 50 mM Tris-HCl (pH 7.5), 200 mM KCl, 2 mM MnCl_2_, 5 mM DTT, and 20% glycerol. 500 μM of dNTPs and 500 nM of reverse primer were added. Reverse transcription reactions were incubated for 3 hours at 42°C, then purified using Ampure XP Beads (Beckman Coulter, A63880). PCR was carried out with target-specific primers and Phusion™ High-Fidelity DNA Polymerase (NEB, M0530L), followed by validation by agarose gel electrophoresis and purification with a DNA C&C Kit (Zymo, D4014) or Ampure XP beads (Beckman Coulter, A63880). Amplicon-EZ sequencing and alignment was then carried out by Azenta on PCR products, with reads containing mutations at each m^1^A site of interest categorized as “modified” for downstream analysis and comparisons. For randomized oligos, occurrences of each nucleotide (A/C/T/G) at 3-mer positions upstream and downstream were counted, separating analyses by “modified” and “unmodified” reads with mutations/no mutations at the site of interest, respectively.

## DATA AVAILABILITY

Raw and processed data files from all high throughput sequencing experiments have been deposited in the NCBI Gene Expression Omnibus and will be released on or before publication. Code for in house analysis pipelines is available on the Nachtergaele lab GitHub repository.

## ACKNOWLEDGMENTS

We would like to thank all members of the Nachtergaele and Gilbert labs, and Christopher King for valuable discussion and feedback, and Fabian Menges for mass spectrometry support. We would also like to thank John Tawil for support with early experiments that led us to this work, and the Yale Centers for Genome Analysis and Research Computing for support with RNA sequencing and analysis. We would like to thank Anna Marie Pyle for the gift of the ultraMarathonRT enzyme used in this work. This work was supported by an NIGMS MIRA award (R35GM146919) and NHGRI R01HG011868 to S.N, and by R01GM101316 and NSF award 2330451 to W.G. . R.S. was supported by T32GM145469 and K.A. was supported by an HHMI Gillian Fellowship. Research reported in this publication was also supported by the National Institute of General Medical Sciences of the National Institutes of Health under award number 1S10OD030363-01A1 (Yale Center for Genome Analysis) and made use of the Chemical and Biophysical Instrumentation Center at Yale University (RRID:SCR_021738; equipment was purchased with funds from Yale University).

