## Supplementary Information for "Investigation of TRMT61B methyltransferase activity on mRNA and its effects on translation"

<u>Supplementary information associated with this manuscript:</u>
Supplementary figures 1-3

Supplementary Table 1. Published TRMT61B methylation targets.

Supplementary Table 2. $m^1A$ -IP DESeq2 analysis.

Supplementary Table 3. $m^1A$ -IP single nucleotide $m^1A$ sites.

Supplementary Table 4. IVT pool single nucleotide $m^1A$ sites.

Supplementary Table 5. Example of uMRT mutation signatures in NanoLuc reporter sequence.

Supplementary Table 6. DESeq2 analysis results from IVT pool and ribosome recruitment scores.

Supplementary Table 7. Primer sequences for reverse transcription and PCR.

Supplementary Table 8. Sequences of oligonucleotides used in DART experiments.

### Supplementary Figure 1

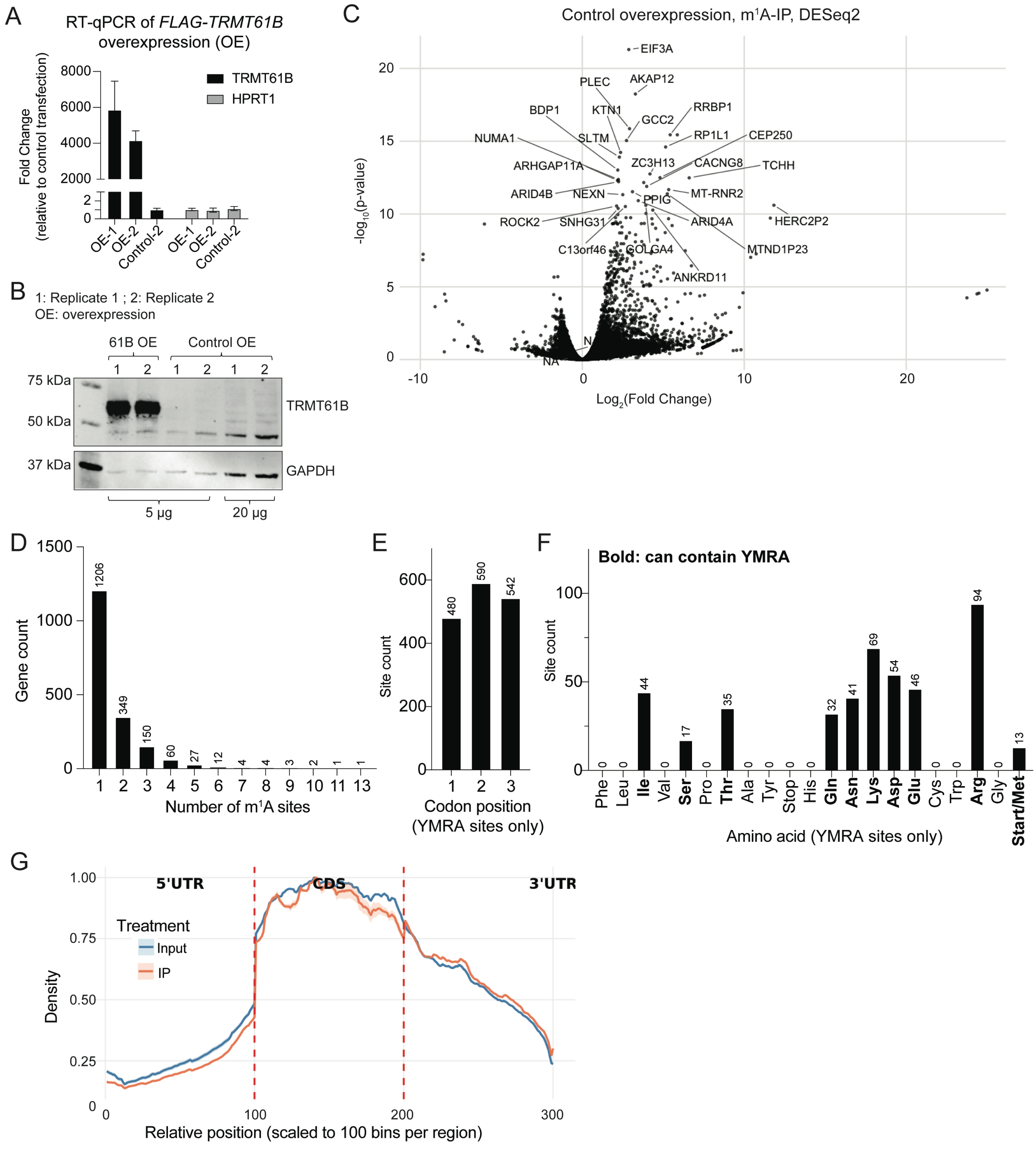


**Supplementary Figure 1. Additional data for TRMT61B overexpression (OE) experiments in U-2 OS cells.** A) RT-qPCR analysis of expression of *TRMT61B* RNA in TRMT61B-FLAG OE and control OE conditions. B) Western blot showing expression of TRMT61B protein in TRMT61B-FLAG OE and control OE conditions. C) Volcano plot showing DESeq2 results depicting fold changes in transcript abundance in m$^1$A-IP samples compared to input in the control overexpression cells. Top 30 hits by -log$_{10}$(p$_{adj}$) are labeled. D) Distribution of the number of high confidence YMRA

$m^1A$ sites per transcript under TRMT61B-FLAG OE conditions. E) Codon positions for high confidence CDS YMRA $m^1A$ sites under TRMT61B-FLAG OE conditions. F) Distribution of the number of high-confidence YMRA $m^1A$ sites from the TRMT61B-FLAG OE $m^1A$-IP found in codons of all possible amino acids. Amino acids with codons that can accommodate a YMRA motif are bolded. G) Metagene analysis of input and IP read coverage across 5'UTR, CDS, and 3'UTR regions. Each region was scaled to 100 bins. Analysis included 5,000 transcripts with all regions ≥50 bp. Coverage was strand-corrected, normalized (0-1 scale), and averaged across transcripts. Lines show the mean of two replicates per condition.

**Supplementary Figure 2**

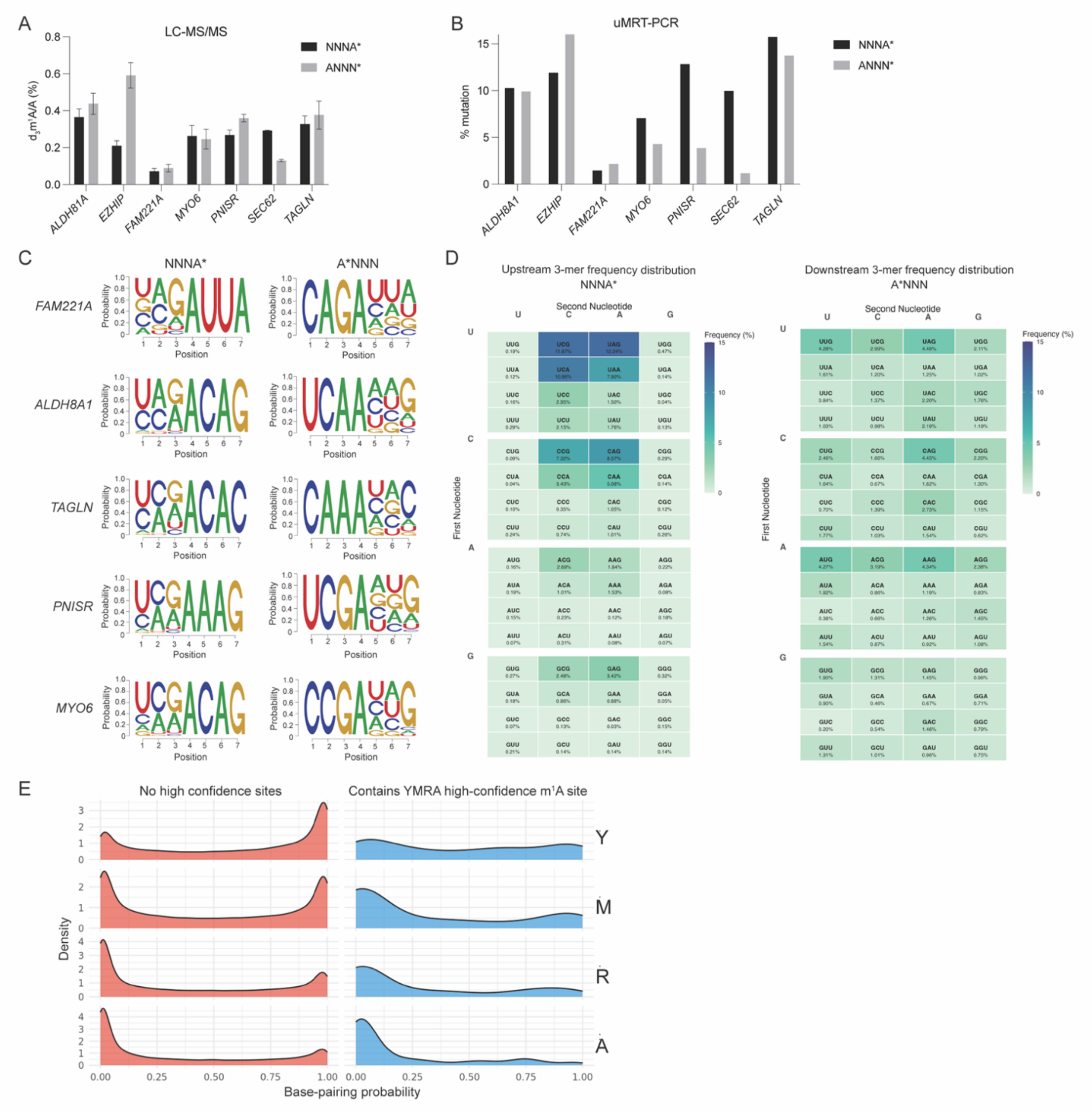


**Supplementary Figure 2. Additional data for randomized motif experiments and m[1]A DART.** A) LC-MS/MS data showing $d_3m^1A$ levels in randomized oligos methylated by TRMT61B-FLAG. B) ultraMarathonRT-PCR data from of randomized oligos methylated by TRMT61B-FLAG, showing misincorporation levels at the specific $m^1A$ modification site of interest. C) Nucleotide abundances at each position between -3 and +3 surrounding the $m^1A$ site of interest, showing only results for "modified" sequences containing a misincorporation at the modification site. D) Frequency of each 3-mer sequence occurring upstream and downstream in "modified" sequences only, combining data from all analyzed randomized sequences. E) Probability of base pairing at each motif position in high confidence methylated YMRA sites (blue) and all other occurrences of the YMRA motif in the 5'UTR/CDS pool (red).

**Supplementary Figure 3**

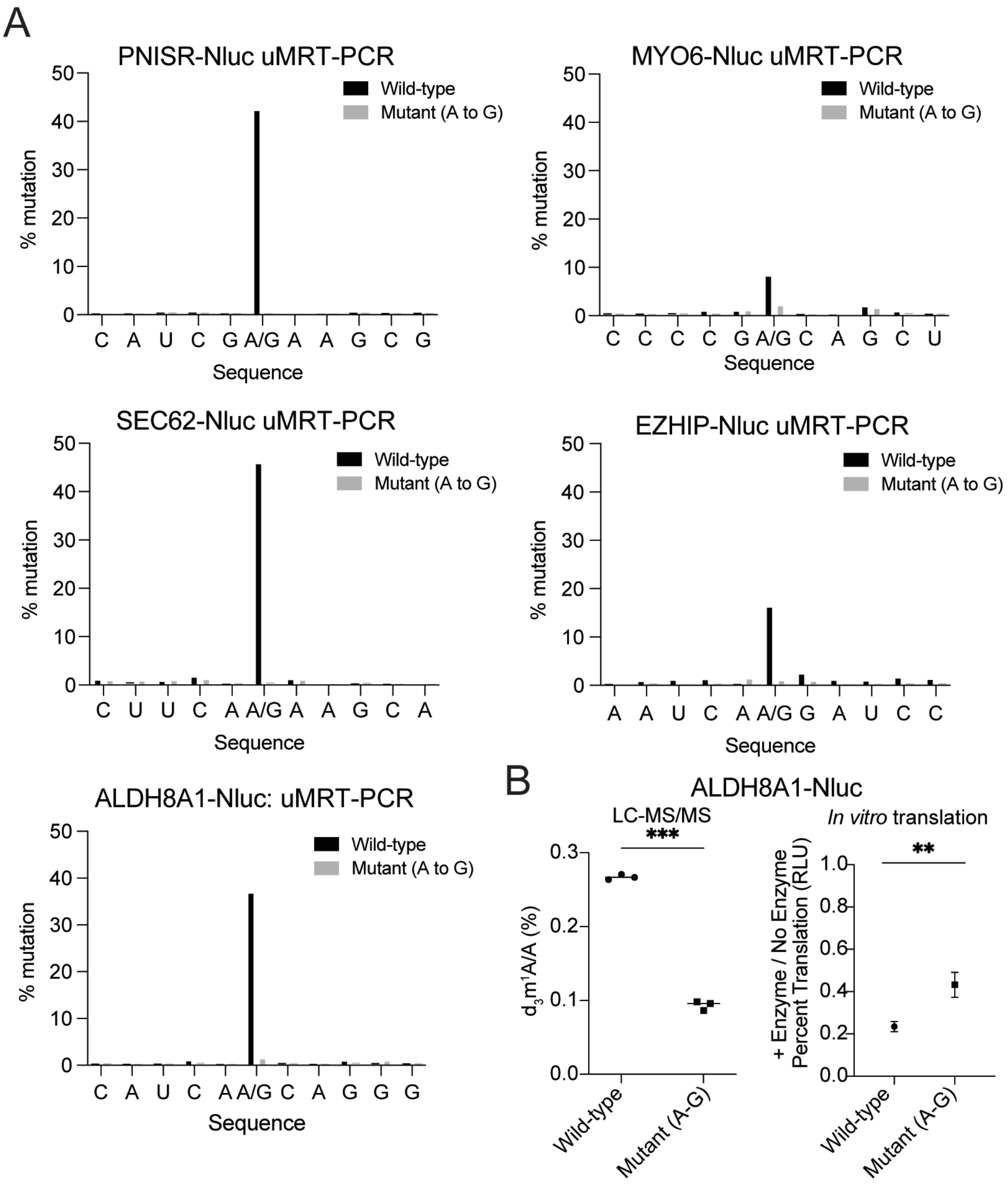


**Supplementary Figure 3. Additional data for *in vitro* luciferase translation experiments.** A) ultraMarathonRT-PCR data for the five tested wild-type and mutant reporter sequences, showing rates of misincorporation following TRMT61B treatment at the site of interest. B) Left: LC-MS/MS measurements of $d_3m^1A$ levels in TRMT61B-FLAG-treated wild-type and mutant ALDH8A1 reporter RNA (additional codon 1 example, related to figure 4). Statistical analysis by unpaired t-test, * p<0.05, ** p <0.01, *** p <0.001, and **** p <0,0001 (n=3 technical replicates). Right: Luciferase translation of TRMT61B-FLAG-treated wild-type or mutant ALDH8A1 reporter RNA, normalized to translation of untreated wild-type or mutant reporter, respectively. Statistical analysis by two-sample z-test comparing independent ratio estimates using propagated standard errors. * p<0.05, ** p <0.01, *** p <0.001, and **** p <0,0001 (n=3 replicate methylation reactions).
